# Neutrophil swarming in damaged tissue is orchestrated by connexin-dependent calcium signals

**DOI:** 10.1101/853093

**Authors:** Hugo Poplimont, Antonios Georgantzoglou, Morgane Boulch, Caroline Coombs, Foteini Papaleonidopoulou, Milka Sarris

## Abstract

Neutrophils are major inflammatory cells that rapidly infiltrate injured tissues to provide antimicrobial functions. A key step in their response is the paracrine release of the attractant LTB4, which switches the migration mode from exploratory patrolling to coordinated swarming. This leads to dense clusters that may further disrupt tissue architecture. The coordination mechanism underpinning neutrophil swarms is elusive. Here we show that neutrophils swarms require mutual reinforcement of damage signalling at the wound core. New biosensors and live imaging in zebrafish revealed that neutrophil chemoattractant synthesis is triggered by a sustained calcium flux upon contact with necrotic tissue and sensing of the damage signal ATP. This ‘calcium alarm’ signal propagates in the nascent neutrophil cluster through connexin-43 hemichannels, which allow release of intracellular ATP. This enables rapid assembly of a centralised, supracellular chemoattractant source, which is instrumental for coordinated recruitment and maximal cell gathering.

## Introduction

Tissue damage triggers rapid recruitment of immune cells, with neutrophils as prime infiltrators (Kolaczkowska and Kubes, 2013; Wang, 2018). This migratory response marks the onset of inflammation, which is essential for protecting the breached tissue from infection while the slow process of tissue repair unfolds. Neutrophils are instrumental for killing bacterial pathogens through phagocytosis, release of proteolytic enzymes and reactive radicals (Kolaczkowska and Kubes, 2013). However prolonged neutrophil residence can cause collateral tissue damage, perpetuate inflammation and delay tissue repair and restoration of homeostasis (Kolaczkowska and Kubes, 2013). Chronic inflammation forms the basis of numerous diseases and can also be co-opted by cancer cells to favour tumor growth and metastasis (Singel and Segal, 2016; Soehnlein et al., 2017). Tuning neutrophil accumulation to desirable levels is thus an important biomedical target, yet our basic understanding of how this response naturally escalates under physiological conditions remains limited.

Interestingly, while the initial steps in neutrophil recruitment are driven by extrinsic cues, the escalation phase of the response is largely self-organised. Tissue injury results in local release of primary damage cues (Damage Associated Molecular Patterns or DAMPs) from necrotic cells, including ATP or formyl peptides, which are normally not present in the extracellular environment (Futosi et al., 2013; Kolaczkowska and Kubes, 2013; Wang, 2018). To a certain extent, these primary signals may act directly as chemoattractants by signalling though corresponding G-protein coupled receptors (GPCRs) (Futosi et al., 2013). Beyond this, DAMPs and other physiological stresses cause secondary production of chemoattractants by local tissue cells, including chemokines or arachidonic acid metabolites (Afonso et al., 2012; Enyedi et al., 2013; Kolaczkowska and Kubes, 2013). Altogether, this cocktail of attractants promotes exit of neutrophils from the blood (extravasation) and biased directional motion (chemotaxis) towards the site of injury within minutes. Thereafter, neutrophil behaviour can switch from mere chemotaxis to highly coordinated and unidirectional motion that culminates into dense clusters at the wound core (Kienle and Lämmermann, 2016; Lämmermann et al., 2013; McDonald et al., 2010). This so-called ‘swarming’ or ‘aggregation’ behaviour is self-organised, as it relies on paracrine release of the lipid attractant leukotriene LTB4 by neutrophils (Afonso et al., 2012; Lämmermann et al., 2013). The decision to release LTB4 is thus critical for the ultimate scale of the response. However, it remains unclear how LTB4 release is triggered and coordinated in individual neutrophils to generate concerted motion.

The dynamics of attractant production have been previously implicated in the reminiscent phenomenon of slime mould aggregation (Dormann et al., 2002). Upon starvation, unicellular amoebae aggregate into a multicellular migratory slug that can seek nutrients. This is driven by initial production of the chemoattractant cAMP in a single amoeba, which triggers further cAMP release in nearby cells, resulting in travelling waves of attractant. Coordination of this response towards a single center-point requires periodic and polarised emission of signal (Dormann et al., 2002; Kriebel et al., 2008). This raises the question whether special mechanisms may have evolved to coordinate neutrophil attractant production and aggregation during tissue damage responses (Afonso et al., 2012; Kienle and Lämmermann, 2016; Sarris and Sixt, 2015). Recent evidence shows that macrophages can prevent swarming by cloaking the wound area, suggesting neutrophil access to the necrotic site is important for swarm decision making (Uderhardt et al., 2019). Another interesting clue is that a critical threshold of initial clustering correlates with subsequent swarming (Park et al., 2018). However, directly relating these observations to chemoattractant production dynamics in neutrophils has been hampered by the lack of tools to monitor the relevant signals directly *in vivo*.

Here we take advantage of the genetic and imaging amenability of zebrafish, in which neutrophil swarming is conserved. We visualised the intracellular events leading to chemoattractant synthesis in individual neutrophils. This revealed that LTB4 is produced in a centralised, collective manner at the wound core by pioneer neutrophils in a nascent cluster, rather than forming spatially autonomous gradients. This is triggered by direct interaction with necrotic tissue as well as mutual reinforcement of damage sensing in clustering cells. Connexin (Cx43) hemichannels, which allow ATP release from live neutrophils, actively amplify damage signalling in the cluster in an autocrine and juxtacrine manner. This communication enables coordinated generation of intracluster calcium fluxes via ATP-gated calcium channels, which in turn promote attractant biosynthesis. Thus, neutrophils cooperate in building a powerful, stable and centralised multicellular gradient source, enabling rapid aggregation at a defined target.

## Results

### Distinct calcium signals in clustering neutrophils

Neutrophil swarming is conserved in zebrafish, a model which is ideally suited for imaging and genetic manipulation (Harvie and Huttenlocher, 2015; Kienle and Lämmermann, 2016). We thus set out to develop tools in zebrafish to study related signalling dynamics. To establish the role of neutrophil LTB4 production in this model, we generated a transgenic zebrafish line, Tg (*lyz:lta4h-*eGFP), expressing leukotriene A4 hydrolase, an enzyme that catalyses the conversion of LTA4 into LTB4 (Peters-Golden et al., 2005), the final step in LTB4 biosynthesis. We used a previously validated translation-blocking morpholino (Vincent et al., 2017) to suppress *lta4h* expression and this led to reduced neutrophil accumulation in wounds (Figure S1A-C). In contrast, *lta4h* knockdown did not affect neutrophil accumulation in wounds of Tg (*lyz:lta4h*-eGFP) larvae (Figure S1D). This confirmed that neutrophil-derived LTB4 drives neutrophil accumulation at wounds, as observed in mammalian systems.

LTB4 production requires calcium-dependent translocation of biosynthetic enzymes to membrane compartments where lipid metabolism takes place (Luo et al., 2003). Intracellular calcium dynamics have been observed in zebrafish epithelial cells (Enyedi et al., 2013) and neutrophils migrating in a solitary manner (Beerman et al., 2015), but not in swarming neutrophils. To characterise this, we generated a transgenic line expressing a sensitive calcium indicator, GCamp6F (Chen et al., 2013), in neutrophils Tg(*lyz:*GCamp6F) (Figure 1A and B). Since swarms are more likely to occur under high neutrophil density (Lämmermann et al., 2013; Park et al., 2018; Reátegui et al., 2017), we visualised the behaviour of neutrophils after acute laser wound injury at a site rich in neutrophils, the caudal hematopoietic tissue (CHT), using two-photon ablation (Figure 1C and Video S1). Within 5 minutes, neutrophils began migrating to the wound in a highly directional and coordinated manner, forming clusters at the wound core by 0.5 hours (Video S1). In small-scale swarms, we could distinguish two waves of recruitment (Video S1, example 2). In larger swarms, the response evolved quickly and one main wave was distinguished (Video S1, example 1).

**Figure 1.**
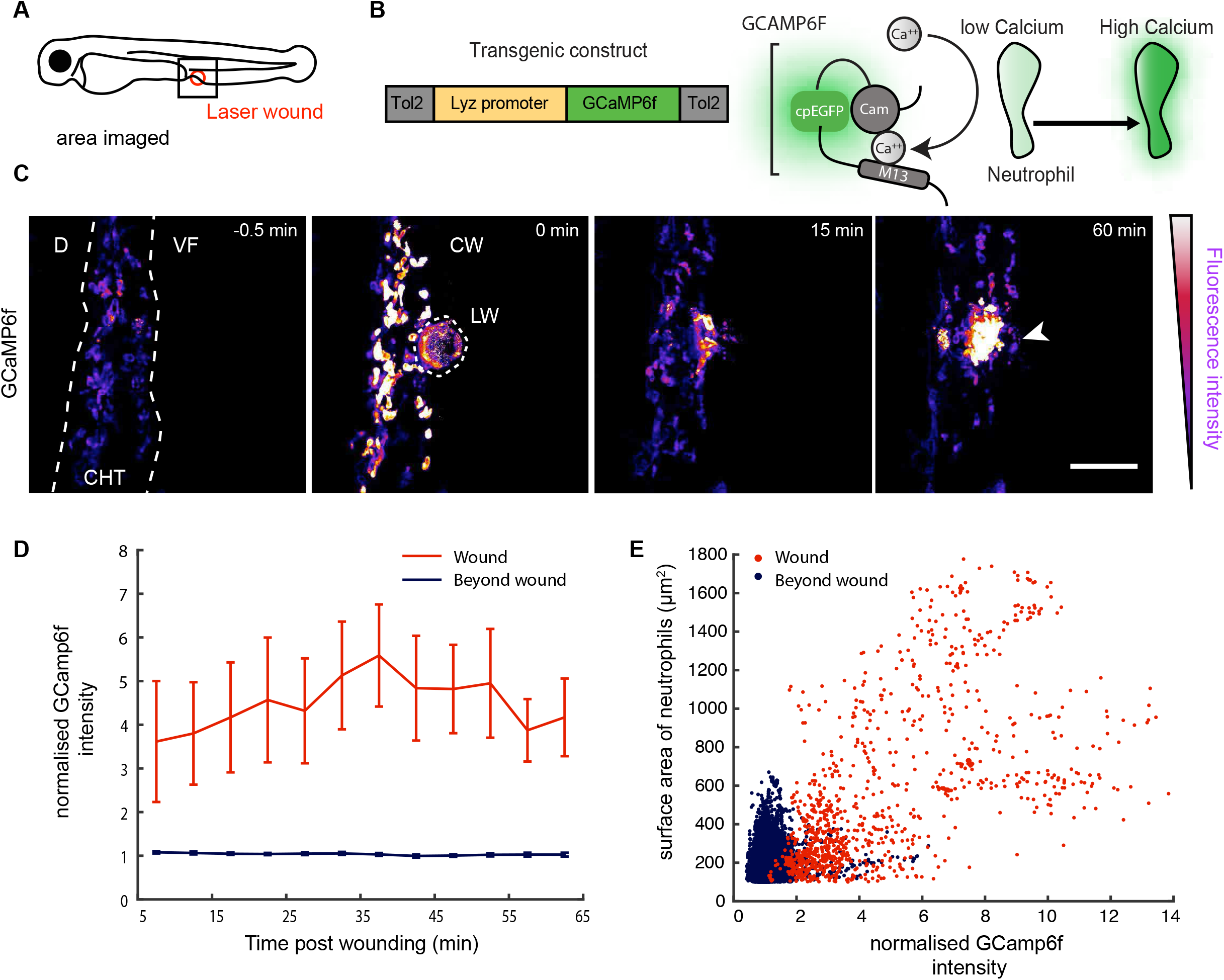
Calcium dynamics in neutrophils during swarming. **A** Schematic of a 3 day-post-fertilisation (dpf) zebrafish larva showing the area of two-photon laser wound damage and imaging. **B** The GCamp6F construct is expressed under the control of the lysozyme C promoter (Lyz). Ca^2+^ binding to the calmodulin (Cam) domain of GCamp6F increases eGFP fluorescence. **C** Time-lapse sequence of neutrophils (colour-coded for GCamp6F intensity) in a Tg(*lyz*:GCamp6F) larva. Images show neutrophils (colour-coded for GCamp6F intensity) within the caudal hematopoietic tissue (CHT) migrating towards a laser wound (LW; dotted line) at the ventral fin-CHT boundary (VF/CHT). The calcium wave (CW) is indicated in the second panel and the neutrophil cluster with a white arrow. Scale bar = 50µm. **D** Quantification of GCamp6F intensity over time in neutrophils at the wound versus beyond the wound (i.e. beyond the laser wound area indicated with dotted line in C). Intensity values were normalised to the mean intensity of segmented neutrophils prior to wound. n=8 larvae in 8 experiments. The first 5 minutes post-wounding are excluded to eliminate the contribution of the calcium wave. **E** Normalised GCamp6F intensity in relation to the surface area of segmented neutrophils. Neutrophils are segmented based on their contour and individual dots represent single neutrophils or clustered neutrophils at the wound (red) or beyond the wound (blue). The more clustered the neutrophils are the larger the surface area detected as an individual neutrophils surface. Indicatively, the surface area for single neutrophils is up to 400 µm^2^. Pooled data from n=8 larvae in 8 experiments.

Regardless of the scale of the response, the intensity of GCamp6F indicated three distinct signals (Figure 1C and Video S1). First, a brief, tissue-wide calcium wave immediately after wounding that dissipated within 30 sec, an anticipated response of tissue to injury (Enyedi et al., 2013; Razzell et al., 2013). Subsequently, neutrophils began migrating and showed dynamic local fluctuations in calcium intensity. In contrast, upon arrival at the wound core, neutrophils underwent a whole-cell, sustained calcium flux concomitant with stopping and clustering (Video S1). Strikingly, the calcium fluxes rapidly propagated across clustering neutrophils giving rise to a cellular mass with relatively sustained calcium signalling (Video S2). Quantification showed that the mean calcium intensity in clustering cells was sustained at high levels throughout the first hour post-wounding (Figure 1D). Moreover, calcium intensity showed a positive association with cluster size (Figure 1E). Altogether this evidence showed that distinct types of calcium signals occur in migrating cells versus clustering cells.

### Activation of LTB4 biosynthesis is restricted to clustering neutrophils with calcium fluxes

The discovery of distinct types of calcium signals in swarming neutrophils prompted us to investigate which of these are consequential on LTB4 biosynthesis. The rate-limiting step in LTB4 biosynthesis is the translocation of 5-lipoxygenase (5-LO or ALOX5) to the nuclear envelope membrane where it converts arachidonic acid into LTA4 (Luo et al., 2003). LTA4 can be further processed to different metabolites, but neutrophils are geared to produce LTB4 (Kienle and Lämmermann, 2016; Serhan and Sheppard, 1990). Thus, 5-LO peri-nuclear translocation provides a microscopically tractable readout to identify neutrophils with activated LTA4/LTB4 biosynthesis (Figure 2A). To link 5-LO translocation with calcium signals, we generated a zebrafish line expressing 5-LO in neutrophils Tg(*lyz*:tRFP-5LO) and crossed this with Tg(*lyz:*GCamp6F) fish (Figure 2B). We monitored 5-LO translocation after acute laser wound, using spinning-disk microscopy. Interestingly, we discovered that peri-nuclear 5-LO translocation occurred principally in neutrophils with a whole-cell calcium flux that were clustering at the wound focus and not in migrating cells (Figure 2C-D, Video S3, example 1). As further evidence, we performed a small incision in the ventral fin, which triggers less compact clusters that facilitate visualisation of subcellular dynamics. We discovered the same trend, in that peri-nuclear 5-LO translocations were detected only among the clustering, calcium-fluxing cells (Figure 2C-D, Video S3, example 2). These data suggested that the specific calcium fluxes observed in clustering cells are associated with activation of LTB4 synthesis.

**Figure 2.**
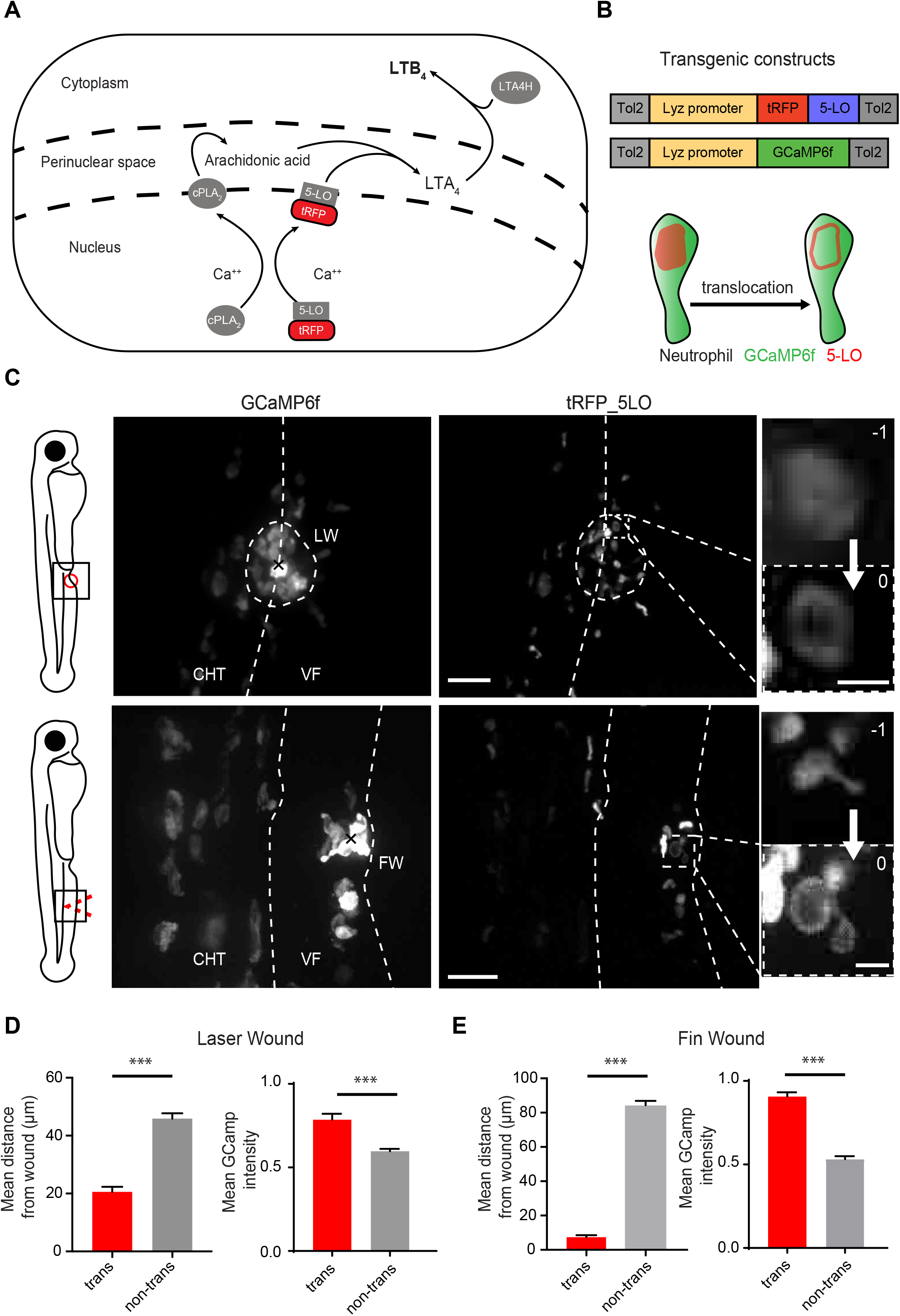
Activation of LTB4 biosynthesis is limited to clustering neutrophils. **A** Schematic of LTB4 biosynthesis. cPLA2 (calcium-dependent phospholipase A2) and 5-LO (5-lipoxygenase) are recruited to the nuclear membrane and produce arachidonic acid (AA) and LTA4 respectively. LTA4 is metabolised into LTB4 by LTA4 hydrolase. **B** Constructs for transgenic expression of a fluorescent fusion of 5-LO with tRFP in neutrophils (below). Schematic of neutrophil with 5-LO nuclear translocation. **C** Time-lapse images of neutrophils in 3dpf double-transgenic Tg(*lyz*:GCamp6F)xTg(*lyz*:tRFP-5LO) zebrafish larvae taken after two-photon laser wound in the ventral fin-CHT boundary (top) or mechanical injury in the ventral fin (bottom). Zoomed images of neutrophils showing 5-LO translocation (time in relation to translocation is indicated in minutes). Scale bars = 50µm and 5µm, respectively. **D-E** Quantification of mean distance from the wound centre (x) and normalised GCamp6F fluorescence intensity for 5-LO-translocating cells versus non-translocating cells in laser wounds **D** and fin wounds **E**. Data are from n=42 translocating and n=720 non-translocating cells (**D** left) and n=31 translocating and 289 non-translocating cells (**D** right) from 8 larvae in 5 different experiments. Data are from n=17 translocating and n=634 non-translocating cells (**E** left) and n=16 translocating and 582 non-translocating cells (**E** right) from 5 larvae in 3 different experiments. Two-tailed unpaired t-test.

### Neutrophil calcium fluxes are triggered upon contact with necrotic tissue or pioneer fluxing neutrophils

Based on these observations, we set out to investigate the mechanism driving the 5-LO-capacitating calcium fluxes in clustering neutrophils. One possibility was that the observed high calcium fluxes resulted from loss of calcium homeostasis due to neutrophil death. Alternatively, the calcium fluxes could have been due to active signalling. To explore these scenarios, we imaged neutrophil wound responses in Tg(*lyz:*GCamp6F) in the continuous presence of propidium iodide (PI) in the bath of the larva. PI is cell impermeable and selectively stains nucleic acids in cells with impaired membrane integrity. The dye cannot penetrate the skin but superficial wounding permitted its transient interstitial access and staining of the local necrotic cells around the wound margin (Figure 3A, Figure S2A and Video S4). Dying neutrophils were distinguished by loss of GCamp6F signal followed by uptake of PI stain (Figure S2B and Video S5). We found that the percentage of these dying neutrophils within the cluster was relatively low (Figure S2C). Moreover, the dying cells ejected themselves on the side of the cluster rather than occupy space in the wound focus (Video S5). Altogether, this suggested that the 5-LO-capacitating calcium signals in clustering cells were unlikely to be mediated in dying neutrophils. Instead, we found that the 5-LO-capacitating calcium fluxes were correlated with necrotic sensing. Specifically, neutrophils underwent a calcium flux concomitant with abrupt deceleration of migration upon direct contact with necrotic cells (Figure 3B and C) or after contact with other pioneer clustering neutrophils at the wound core (Figure 3D and Video S2).

**Figure 3.**
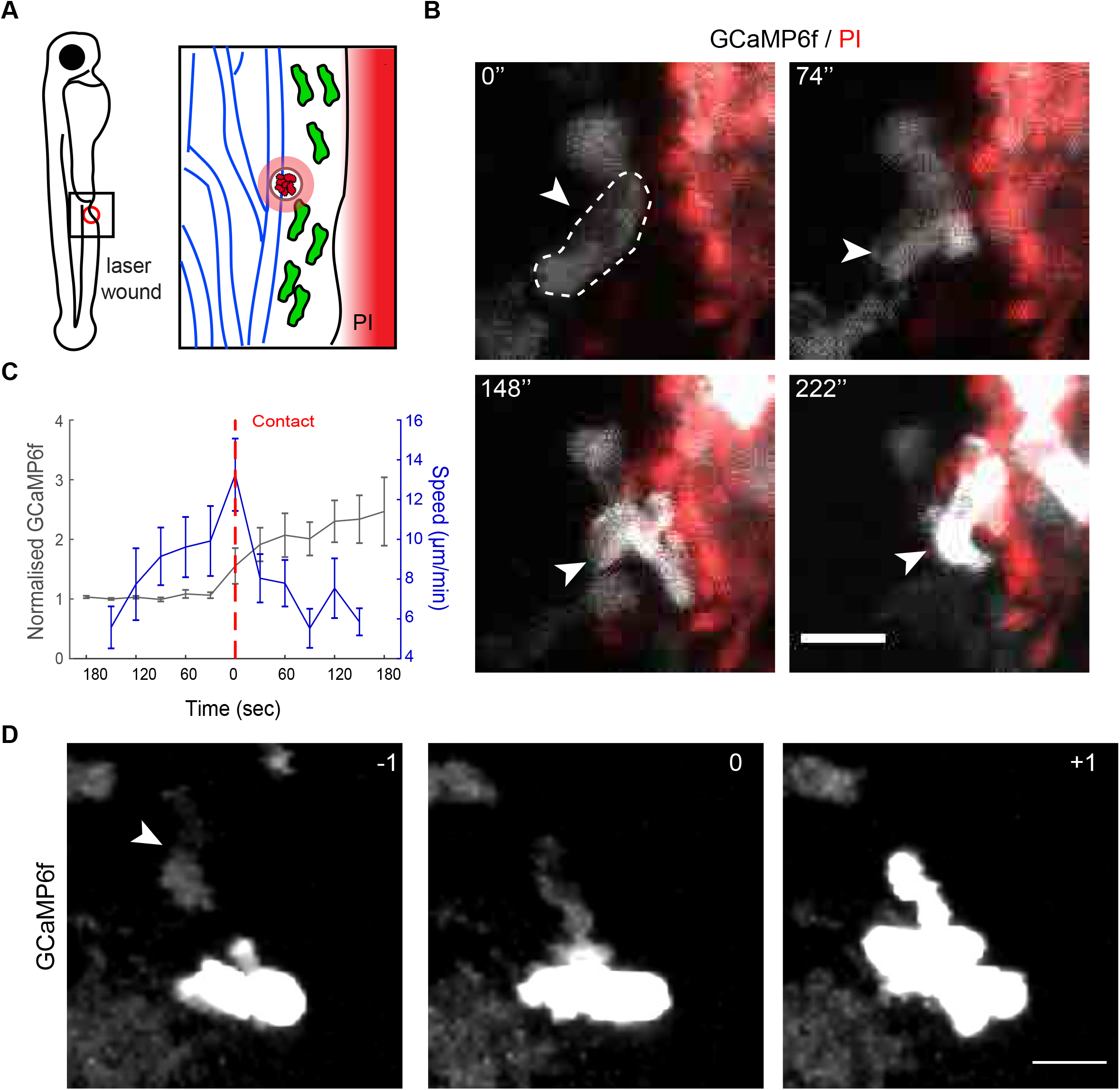
5-LO-capacitating calcium fluxes are triggered upon contact with necrotic cells or neutrophils with ongoing fluxes. **A** Schematic of two-photon laser wounding on 3dpf larvae in the continuous presence of propidium iodide (PI) in the embryo medium. **B** Time-lapse images of a GCamp6F-expressing (white) neutrophil (indicated with an arrow) entering a contact with PI^+^ tissue cells (red); Time in relation to the first frame is indicated in seconds. Scale bar = 10µm. **C** Quantification of speed (blue) and normalised GCamp6F (grey) in neutrophils before and after contact with PI^+^ tissue. Dotted red line indicates time of contact. Pooled cell data n=23 cells from 7 larvae in 4 experiments. **D** Time lapse images of a dim GCamp6F^+^ neutrophil (arrow) contacting a bright GCamp6F^+^ neutrophil. Time in minutes is indicated relative to cell-cell contact. Scale bar = 10 µm. Quantification of such contacts is shown in Figure 5 G.

### Extracellular calcium entry through ATP-gated calcium channels promotes activation of LTB4 biosynthesis

We then investigated which necrotic tissue signal might elicit 5-LO-capacitating calcium signals in neutrophils. A calcium signal of reminiscent amplitude has been observed in human neutrophils *in vitro* upon activation of the ATP-gated calcium channel P2X1 (Wang et al., 2017). To address whether this pathway was responsible for the calcium fluxes in clustering neutrophils *in vivo*, we used an inhibitor for P2X1, NF279. We found that this inhibitor markedly suppressed the calcium fluxes and 5-LO perinuclear translocation in clustering neutrophils (Figure 4A-E and Video S6). We then tested whether secondary chemoattractants, such as chemokines, would be able to trigger a sustained calcium flux. We monitored neutrophil behaviour and calcium dynamics in the presence of Cxcl8a-mCherry secreting transplanted cells, which we have previously shown to form extracellular chemokine gradients in vivo (Sarris et al., 2012). Neutrophils accumulated in the transplant but did not show whole-cell calcium fluxes (Video S7 and Figure S3).

**Figure 4.**
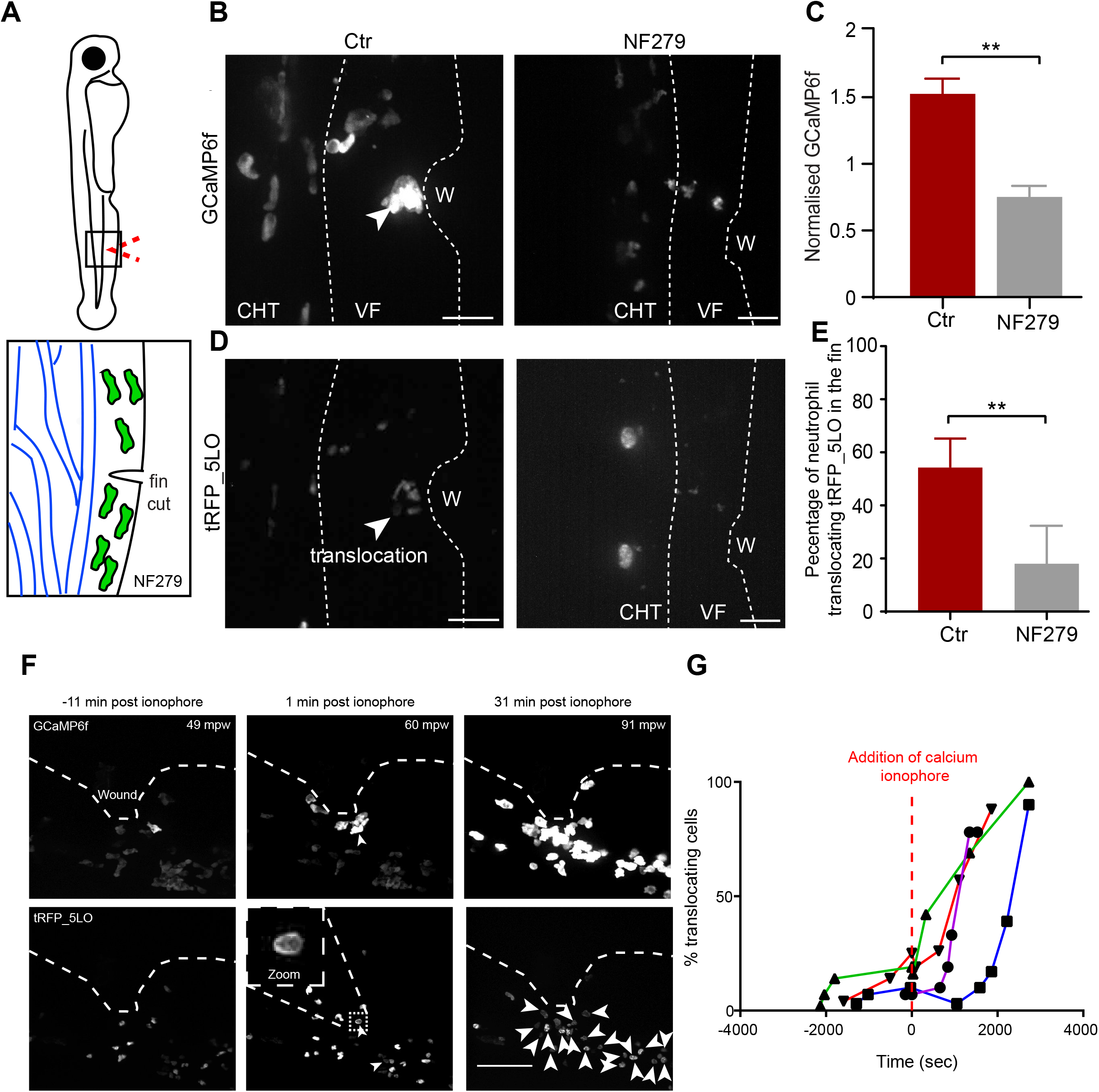
ATP-gated calcium channels and extracellular calcium entry trigger 5-LO-capacitating calcium fluxes in neutrophils *in vivo*. **A** Schematic of ventral fin wounding in the presence of NF279. **B** Maximum intensity projection images of neutrophils in Tg(*lyz*:GCamp6F)xTg(*lyz*:tRFP-5LO) zebrafish larvae 120 minutes after ventral fin wound (VF) in the presence (right) or absence (left) of 10µM NF279. Scale bar = 25µm. **C** Mean normalised GCamp6F intensity in larvae with and without NF279. NF279-treated larvae: n= 3 larvae; control larvae; n=9 larvae. Mann-Whitney test. **D** Maximum intensity projection images of neutrophils in Tg(*lyz*:GCamp6F)xTg(*lyz*:tRFP-5LO) zebrafish larvae 120 minutes after mechanical ventral fin wound (VF) in the presence (right) or absence (left) of 10µM NF279. Scale bar = 25µm. **E** Percent of translocating neutrophils out of all neutrophils recruited into the ventral fin over two hours of imaging starting 10 min post-wounding. Data are means of n=7 control larvae and n=4 NF279-treatd larvae. Mann-Whitney test, two tailed, p-value= 0.0030. **F** 3dpf zebrafish larvae Tg(*lyz*:Gcamp6f) were wounded on the ventral fin (VF) and subsequently imaged on a spinning disc microscope for up to 120 minutes post-wounding. 50µM calcium ionophore (A23187) was added after migration initiation, typically between 30- and 60-minutes post-wounding (mpw). The top images show confocal projections of the Gcamp6f channel and the bottom images show the 5-LO-tRFP channel at the indicated time points after wounding (the time relative to calcium ionophore is shown above the images). Arrows indicate translocation events. Scale bar = 50µm. **G** Percent of 5-LO tranlocating neutrophils out of all neutrophils visible in the field of view over time. The time of calcium ionophore addition is set to 0. n=4 larvae from 2 different experiments. Each line represents an individual embryo.

These data suggested that the prominent calcium fluxes of clustering neutrophils at the wound core act as an alarm system triggered by calcium channel opening upon exposure to a threshold of the damage signal ATP (we hereafter refer to this as ‘calcium alarm signal’). To establish whether extracellular calcium entry is sufficient to trigger 5-LO translocation and stopping, we added a calcium ionophore after neutrophils initiated migration to the wound. Within minutes, neutrophils experienced a calcium flux that led to immediate arrest and concomitant 5-LO translocation (Figure 4F, G and Video S8).

### Neutrophil connexin-43 is required for coordinated calcium fluxes and swarm initiation

We had so far identified ATP sensing as a primary trigger for the calcium fluxes underpinning neutrophil swarming. However, this did not explain the highly efficient and coordinated spread of calcium fluxes within clusters. We hypothesised that clustering neutrophils may be mutually reinforcing ATP signalling. *In vitro* studies suggest that neutrophils can release ATP through connexin hemichannels but the physiological relevance of this finding has not been explored *in vivo* (Bao et al., 2014; Eltzschig et al., 2006). To test whether connexin channels contribute to neutrophil calcium alarm signals *in vivo*, we visualised neutrophil behaviour in the presence of carbenoxolone (CBX), a drug that inhibits connexin channel activity (De Vuyst et al., 2007). This treatment profoundly inhibited neutrophil calcium alarm signals and led to exploratory single cell motility (Figure 5A and B and Video S9). To quantify this, we measured the radial speed of neutrophils over time, which reflects the level of coordination of migration (Lämmermann et al., 2013). When cells move in synchrony in the same direction, the amplitude of radial speed of the population is high. Accordingly, in non-treated neutrophils, we detected a marked wave of synchronous directional motion, peaking at 15 min post-wounding. This time course is comparable with the evolution of radial speed in mouse neutrophil swarms (Lämmermann et al., 2013). We found that CBX treatment led to a marked reduction in the amplitude of the wave of neutrophil migration, consistent with loss in coordination (Figure 5C). To genetically corroborate this, we investigated connexin expression in zebrafish neutrophils and found two connexin genes to be expressed, *cx43* and *cx43.4* (Figure S4A and B). We found that combinatorial CRISPR/Cas9 knockout led to zebrafish embryonic lethality before the onset of neutrophil development. This is consistent with the embryonic lethality of Cx43 null mutations observed in mice (Lo et al., 1999). We thus used knockdown with *cx43* morpholinos that phenocopy hypomorphic mutations of *cx43* (Hoptak-Solga et al., 2008; Iovine et al., 2005). The morpholino mixture for *cx43*/*cx43.4* resulted in reduced retina size, an expected developmental phenotype (Iovine et al., 2005) (Figure S4C). Consistent with the CBX results, Cx43 knockdown reduced neutrophil calcium fluxes and coordination of migration (Figure 5A-C and Video S9). Importantly, both the CBX treatment and the Cx43 knockdown severely compromised the transmission of calcium fluxes across contacting neutrophils (Figure 5F-H).

**Figure 5.**
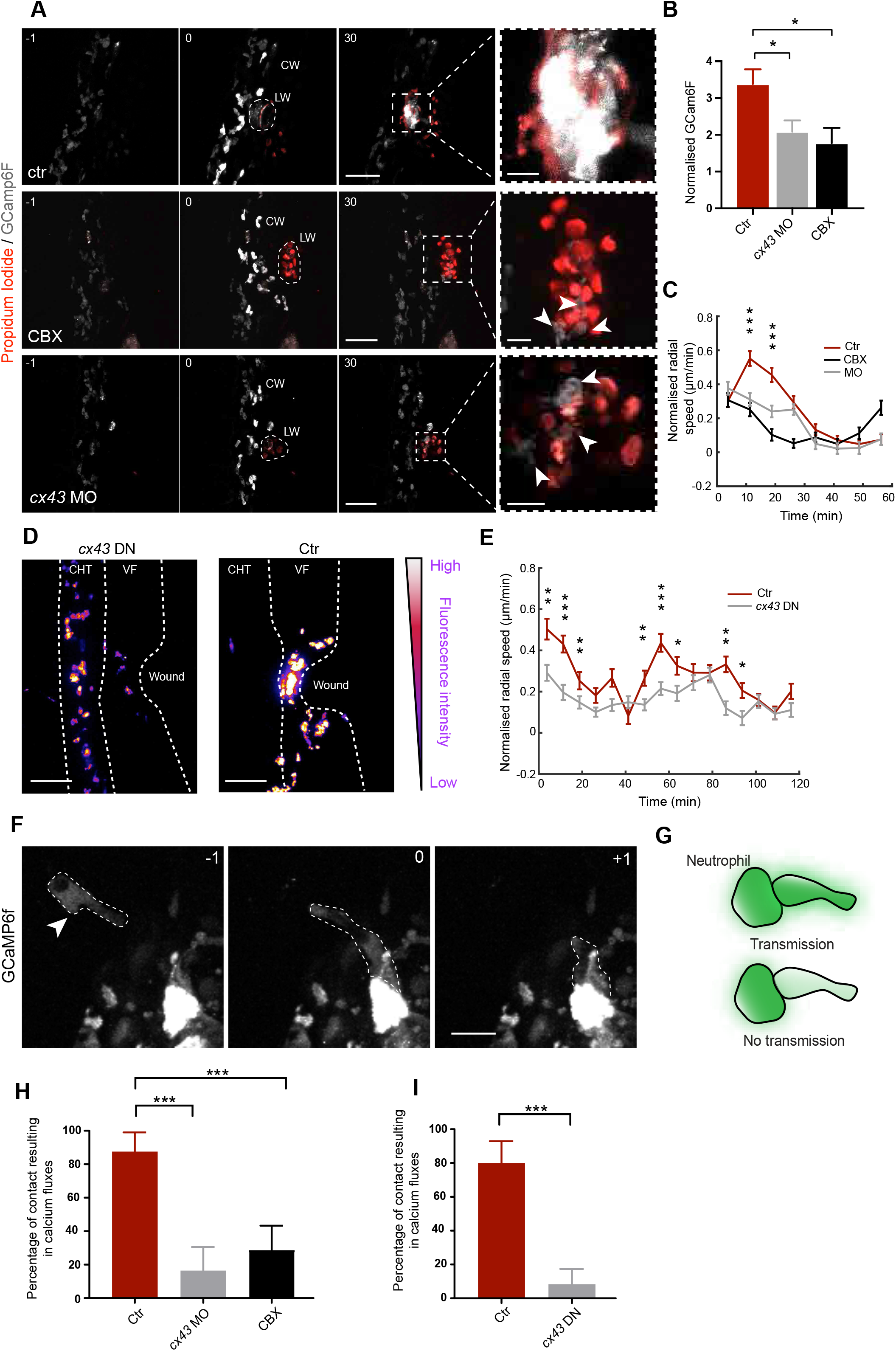
Neutrophil Cx43 is required for coordinated calcium fluxes and swarm initiation. **A** Neutrophils in Tg(*lyz*:GCamp6F) larvae in the presence of PI, without treatment (ctr), with 50 µM CBX or with morpholinos against *cx43*/*cx43.4* (*cx43* MO). Scale bars = 50µm and 10µm, respectively. CW; calcium wave. Time after laser wound (LW) is shown in minutes. **B** Normalised GCamp6F levels in control, *cx43* MO-treated and CBX-treated larvae. n=8 control, n=7 *cx43* MO-treated, n=5 CBX-treated from 8, 3 and 2 experiments respectively. One-way ANOVA with Dunnet’s post-test. **C** Normalised radial speed-time plots of neutrophils. Instantaneous speed values for individual neutrophils were divided by the mean instantaneous speed value of the corresponding embryo to normalize differences in speed across embryos (see also methods). Each line shows pooled cell data from multiple larvae binned every 7.5 min. Pooled cell data from control (n=12), CBX-treated (n=8) or MO-injected larvae from x-y experiments. Mann-Whitney test results between ctr and MO shown. **D** Neutrophils in Tg(*lyz*:GCamp6F)xTg(*lyz*:*cx43dn-*T2A-mCherry) zebrafish larvae, positive (*cx43* DN) or negative for the transgene (Ctr), 2h after mechanical fin wound. GCamp6F channel is shown, colour-coded for intensity. **E** Normalised radial speed-time plots of neutrophils. Data pooled from n=11 *cx43DN* transgenics and n=8 control siblings from 4 experiments. Mann-Whitney test. **F** Time lapse images of a dim GCamp6F^+^ neutrophil (arrow) contacting a bright GCamp6F^+^ neutrophil in a CBX treated larvae. Time in minutes is indicated relative to cell-cell contact. scale bar = 15µm. **G** Cartoon shows two types of contacts counted in **H** and **I** between bright and dim Gcamp6F+ neutrophils. **H** Quantification of the percentage of neutrophil contacts resulting in transmission of calcium fluxes. Contacts in which none of the cells is initially fluxing were not included in this analysis. N = 8 control, 7 *Cx43* morphants and 4 CBX-treated larvae from 8, 3 and 2 experiments respectively. One-way ANOVA, Tukey’s multiple comparisons test. **I** Percentage of neutrophil contacts resulting in transmission of calcium fluxes. n=11 *cx43DN* transgenics and n=8 control siblings from 4 experiments. Mann-Whitney test.

Given the broad expression of *cx43*, we next interrogated whether neutrophil *cx43* is important for neutrophil swarming. To this end, we generated transgenic zebrafish whereby neutrophils express a dominant-negative version of *cx43* (Tg (*lyz:cx43DN*-T2A-mCherry) (Oyamada et al., 2002). The behaviour of neutrophils in these transgenics was similar as in *cx43* morphants, in that they showed reduced whole-cell calcium fluxes and clustering at the wound core and less coordinated motility (Figure 5D-E and Video S10). Moreover, we did not observe contact-dependent propagation of calcium fluxes under Cx43 inhibition (Video S11 and Figure 5I). Assessment of neutrophil accumulation at fixed time points across a large pool of embryos showed that inhibition of neutrophil Cx43 suppressed neutrophil accumulation to a similar degree as global Cx43 inhibition (Figure S5). This suggested that neutrophil Cx43 largely accounts for the overall defect in neutrophil accumulation at wounds. Altogether, this evidence demonstrated an important role for neutrophil Cx43/Cx43.4 in coordinating neutrophil calcium signalling and swarming.

Cx43 has channel-dependent and channel-independent functions (Kotini et al., 2018; Ribeiro-Rodrigues et al., 2017). CBX and Cx43DN act on connexin channel activity, suggesting a channel-dependent function in this context. However, it remained unclear whether connexins mediate gap-junction coupling or hemichannel-based signal transmission in neutrophils. We did not observe PI uptake in live neutrophils, suggesting either absence of hemichannel activity or that the level of transport is below our detection limit (hemichannel opening would presumably be transient and brief unlike the permanent membrane integrity disruption in necrotic cells). We thus used functional tests to interrogate a link between Cx43 and P2X1 signalling in neutrophil swarming *in vivo*. Specifically, we found that Cx43 inhibition did not cause further reduction in neutrophil accumulation in NF279-treated larvae (Figure S6). This indicated that Cx43 and P2X1 act in the same pathway, providing evidence for hemichannel-based relay of ATP signalling.

## Discussion

Neutrophil accumulation in inflamed tissue has pervasive implications in disease and therapeutic strategies to fine-tune this process are desirable. However, an understanding of how this migratory response physiologically escalates is lacking. Here, we reveal a cascade of signalling events that underpins amplification of neutrophil migration into prominent swarms. Our experiments reveal that Cx43 hemichannels drive purinergic activation of calcium fluxes in a primary cluster of neutrophils, which in turn fuels mass production of the attractant LTB4 in the growing aggregate. As Cx43 hemichannels mediate ATP release from live cells (Eltzschig et al., 2006; Wang et al., 2017), we propose that neutrophil Cx43 hemichannels actively amplify damage sensing in an autocrine and juxtacrine fashion at the wound centre, such that a centralised, powerful gradient source of LTB4 can be assembled by a core group of cells (Figure 6).

**Figure 6:**
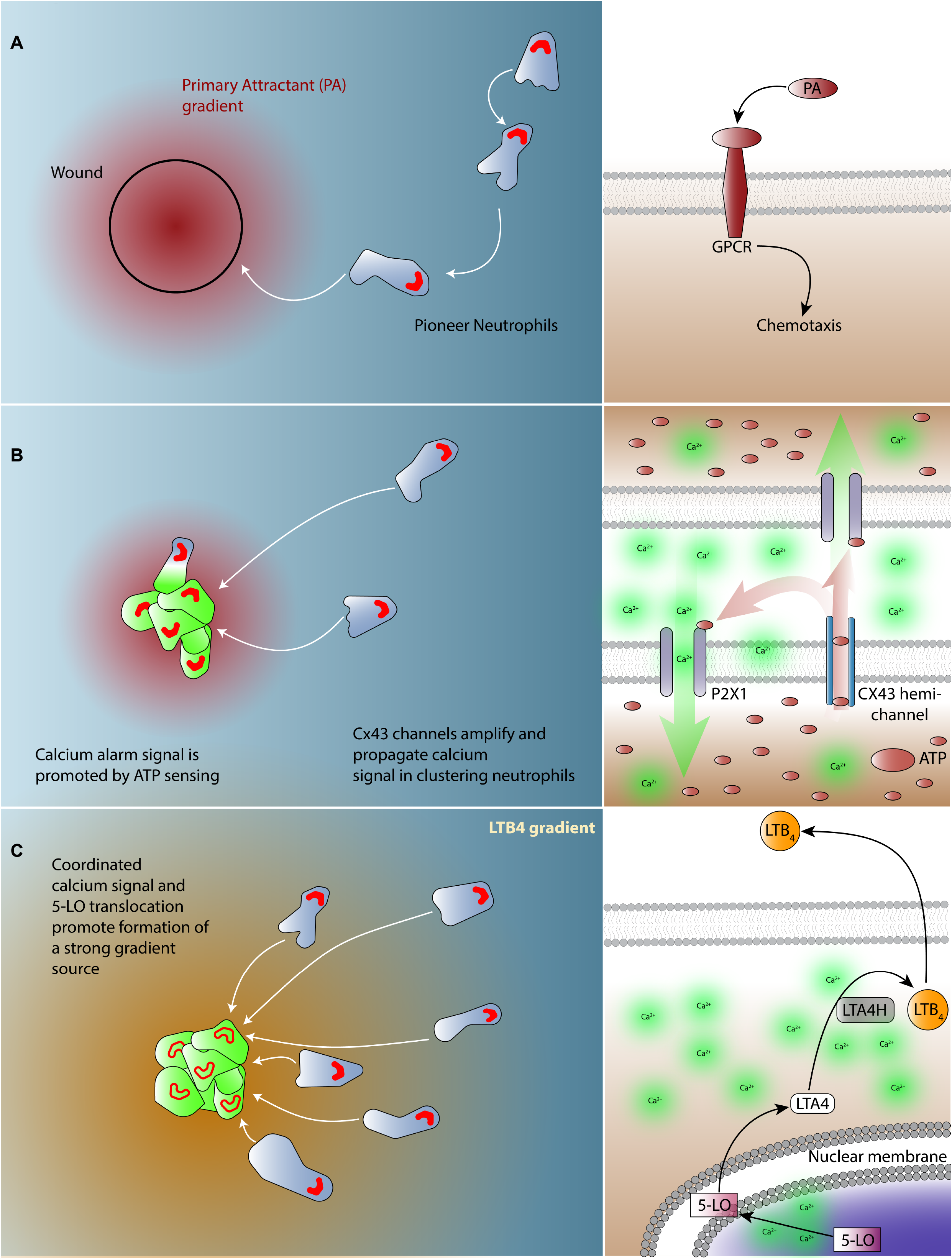
Model for initiation of neutrophil swarms by connexin-dependent calcium signals. **A** Pioneer neutrophils are attracted by primary attractants/cues (DAMPs) to the wound site. The distribution of 5-LO and levels of calcium are indicated in red and green respectively. **B** The primary attractant gradient may decay but pioneer neutrophils are triggered to release ATP through Cx43 channels, amplifying damage sensing in an autocrine and juxtacrine manner. This prolongs the half-life of damage signals, ‘buying time’ for the formation of an effective multicellular LTB4 gradient. Threshold levels of ATP lead to opening of P2X1 channels and a rise of intracellular calcium that leads to neutrophil arrest and 5-LO translocation, ultimately activating LTA4/LTB4 synthesis and further clustering. Note that the intracellular concentration of ATP is high compared to extracellular levels, and the opposite gradient applies in the case of calcium. **C** Coordinated activation of LTA4/LTB4 synthesis in the cluster progressively builds a powerful and stable LTB4 gradient source that triggers a coordinated wave of migration.

The mechanism we describe explains several observations, such as why neutrophil access to the necrotic site is crucial for initiation of swarming (Uderhardt et al., 2019) and why a primary neutrophil cluster precedes the onset of rapid aggregation (Chtanova et al., 2008; Lämmermann et al., 2013; Park et al., 2018). Our model also provides a plausible explanation for the far reaching radius of LTB4-driven chemotaxis during swarming (Lämmermann et al., 2013), as the theoretical range of a gradient is dependent on the concentration of signal produced at the source and its diffusion and degradation rate (Crick, 1970). The assembly of a substantial, multicellular attractant source combined with the rapid tissue diffusion of LTB4 (Demy et al., 2017) could therefore increase the radius of the corresponding chemical gradient. This does not exclude the contribution of LTB4-containing exosomes in the process, whose slower release and propagation could have additional delayed effects in the response (Lim et al., 2015; Majumdar et al., 2016). Our neutrophil-specific inhibition of Cx43 revealed the role for these channels for intracluster calcium alarm signals and swarm initiation. This could provide an explanation for the observed reduced neutrophil accumulation in mouse wounds of Cx43-deficient mice, a phenotype previously presumed to involve endothelial Cx43 and neutrophil extravasation defects (Qiu et al., 2003).

The coordination mechanism we describe is distinct from previous paradigms of collective cell migration. In a cohesive migrating group of cells, such as neural crest cells or the lateral line primordium, intercellular adhesion is critical for ensuring coordination of motion (Scarpa and Mayor, 2016). Cohesiveness further allows self-determination of directionality through asymmetric distribution of receptors across the moving cell mass (Donà et al., 2013; Venkiteswaran et al., 2013). In the non-cohesive paradigm of slime mould aggregation, pulsatile release of attractant underpins coordinated gathering towards a single cell, the location of which is arbitrary (Dormann et al., 2002). Here, we show that intercellular signal amplification within a seeding cluster powers the formation of a critically strong attractant source. This simple mechanistic principle appears to balance the benefit of rapid escalation with the risk of excess or mistargeted congregation. The requirement for close cooperation in the primary cluster provides a level of stringency in the initiation of swarming. The focalised activation of attractant biosynthesis at the wound core provides spatial restriction.

Our study points to interesting future lines of inquiry. One issue that remains unclear is why the activation of 5-LO capacitating calcium fluxes is restricted to the clustering cells at the wound core. P2X1 channel opening requires a threshold level of ATP and such levels might be more likely encountered at the wound core if connexin hemichannels are selectively activated in this locus. Dying tissue cells or dying neutrophils at the wound or specialised pioneer neutrophils could be releasing signals that activate connexin hemichannel opening. These could include LTB4 or fMLP (Eltzschig et al., 2006). We envisage that a high threshold of these molecules might be required for hemichannel activation, given the lack of 5-LO activation in migrating cells distant from the wound. Interestingly, mechanical stimulation of the nucleus plays a role in 5-LO activation (Enyedi et al., 2016) and this may be a cofactor in spatially restricting the source. Another key point to elucidate will be the factors that terminate neutrophil swarming. Based on our imaging, the duration of 5-LO translocation is difficult to track in the dense clusters but it is noteworthy that the relevant calcium fluxes appear to be sustained beyond the duration of the migration wave. As monocyte/macrophage recruitment correlates with cessation of swarms (Lämmermann et al., 2013), it would be interesting to explore how these cells may modulate neutrophil-derived metabolites in this context.

Altogether, our study describes a novel mechanistic paradigm of collective cell behaviour and identifies connexin channels as a key determinant of neutrophil swarming. This opens several avenues for investigation of this pathway in physiological and pathological conditions.

## Acknowledgments

We thank Philippe Bousso, Menna Clatworthy, Ewa Paluch and Rob White for comments on the manuscript, Kevin O’ Holleran for two-photon microscopy, Bill Harris and Christine Holt groups for confocal microscopy, Nachiket Kashikar for the GCamp6F cDNA, Anna Huttenlocher for the *lyz* backbone vector, J.P. Levraud for cDNA of adult zebrafish, Steve Renshaw for the Tg (*mpx*:GFP)^i114^ line. H.P. was supported by a Wellcome Trust PhD grant (105391/Z/14/Z). M.S. and the research was supported by an MRC CDA (MR/L019523/1), a Wellcome Trust [204845/Z/16/Z]; Isaac Newton Trust [12.21 (a)i] and a Royal Society Research Grant (RG170247). M. Boulch was supported by an Erasmus programme (Master de Biologie, École Normale Supérieure de Lyon). C. Coombs was supported by an MRC DTP programme. F. Papaleonidopoulou was supported by an Erasmus programme.

The authors declare no competing interests.

## Methods

### General zebrafish procedures

Zebrafish were maintained in accordance with UK Home Office regulations, UK Animals (Scientific Procedures) Act 1986. Adult zebrafish were maintained under project licence 70/8255, which was reviewed by the University Biomedical Services Committee. Animals were maintained according to ARRIVE guidelines. Zebrafish were bred and maintained under standard conditions at 28.5 ±0.5°C on a 14h light: 10h dark cycle. Embryos were collected from natural spawnings at 4-5 hours post-fertilization (hpf) and thereafter kept in a temperature controlled incubator at 28°C. Embryos were grown at 28°C in E3 medium, bleached as described in the Zebrafish Book (Westerfield M, 2007) and then kept in E3 medium supplemented with 0.3 µg/ml of methylene blue and 0.003% 1-phenyl-2-thiourea (Sigma-Aldrich) to prevent melanin synthesis. For live-imaging of neutrophils expressing fluorescent receptors, methylene blue was omitted from E3 medium to minimise tissue autofluorescence. All embryos were used between 2.5-3.5 dpf thus before the onset of independent feeding. For live imaging or fixation, larvae were anaesthetised in E3 containing 0.04% MS-222 (Sigma). Where indicated, larvae were treated with 50µM Calcium ionophore A23187 (Sigma), 10µM NF279 (BIO-TECHNE LTD), 50µM Carbenoxolone (Sigma) in E3.

### DNA constructs and transgenic zebrafish lines

All DNA expression vectors for transgenesis use a backbone vector with a Lysozyme C promoter (*lyz*) for neutrophil-specific expression and minimal Tol2 elements for efficient integration and a SV40 polyadenylation sequence (Yoo et al., 2012). The sequences cloned in this backbone vector are represented on table 1.

**Table 1.** Constructs cloned.

| Name | Sequences | Origin |
| --- | --- | --- |
| *gcamp6f* | Chen et al. 2013 | Chen et al. 2013 (Chen et al., 2013) |
| *lta4h-egfp* | *lta4h* cDNA:<br>ENSDART00000028171.7 | Gene synthesised<br>(Genewiz) |
| *trfp-5lo* | *5lo/alox5a* cDNA:<br>ENSDART00000079884.6 | cDNA library from whole adult zebrafish |
| *cx43dn_t2a_mCherry* | Omayada et al. 2002<br>(Oyamada et al., 2002) | Gene synthesised (Genewiz) |

The sequence of zebrafish *alox5/5-lo* was chosen over 4 alox genes on the basis of similarity with human 5-LO (Adel et al., 2016). For transgenesis, 0.5nL of solution containing 25ng/µL DNA plasmid and 35 ng/µL were injected into the cytoplasm of one-cell stage embryos. Transposase mRNA was synthesised from pCS2-TP (citation Tol2kit) by *in vitro* transcription (SP6 message machine, Ambion). Injected embryos were stored at 28°C until 5dpf and thereafter were raised in the fish nursery according to standard rearing protocols. At 3 months old, F0 fish were outcrossed to a wildtype (TL) line in order to screen for germline transgenesis.

### Morpholino knock down

Morpholinos were produced by GeneTools LTD and their names, sequences, types and origins are indicated in table 2. All morpholinos were injected into one-cell stage embryos in a morpholino injection solution (120mM KCl, 20mM HEPES, 0.1% phenol red). We used morpholinos against the two isoforms found in neutrophils: *cx43* (also called *cx43.3*)(Hoptak-Solga et al., 2008; Iovine et al., 2005) and *cx43.4*. 1nL of 0.2mM of each morpholino was injected (0.4mM total). As injection control we used 0.4mM of Negative Vivo-Morpholino control oligo. For *lta4h* knockdown 3nL of 0.5mM *lta4h* MO was injected (Vincent et al., 2017).

**Table 2.**
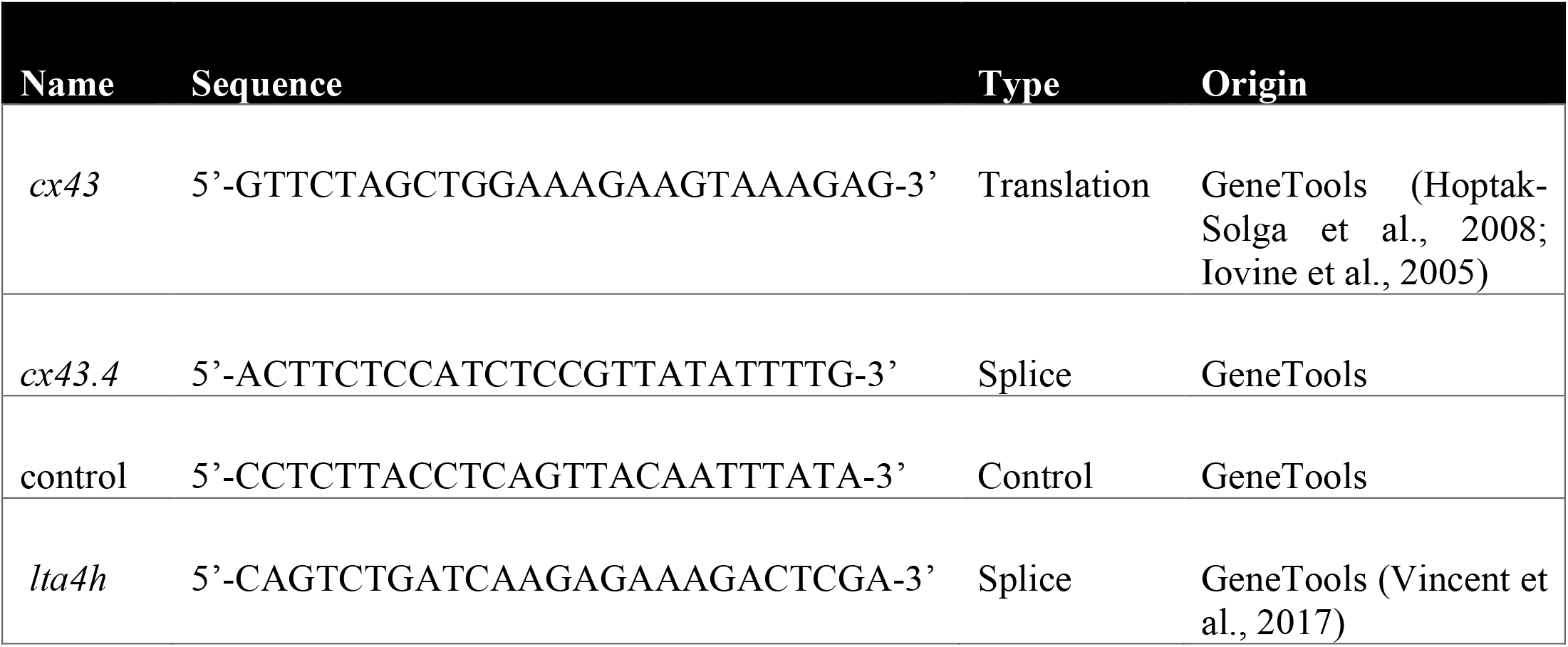
List of morpholinos injected.

### Western blotting

For western blotting, 10 larvae (3dpf) of each genotype were collected. Larvae were then lysed in 100µL of OCG buffer (0.3M NaCl, 2.5µM EDTA pH8, 0.9M Tris HCl pH7.5, protease inhibitors, phosphatase inhibitor) with 1mm glass beads (BioSpec) for 3×20sec in the sonicator Bioruptor^®^ (diagenode). Qubit protein assay kit (Invitrogen) was used to obtain protein concentration. Proteins (25µg) were resolved on a Bolt 10% Bis-Tris Plus Gel (Invitrogen), blotted onto nitrocellulose membrane using iBlot2 transfer stacks (Life Technologies) according to manufacturer’s protocol. Proteins were probed with rabbit anti-human Cx43 (1:2000) (Sigma-Aldrich) and rabbit anti-ß-Tubulin antibodies (1:2000) (Abcam) after saturation in PBST (PBS, 0.1% Tween-20) containing 5% of milk. Proteins were then revealed using an enhanced chemiluminescence detection system (Pierce ECL Plus Western Blotting Substrate, Invitrogen) with goat anti-rabbit HRP antibody (1:2000) (Abcam).

### Two-photon laser wound and live imaging

For mechanical ventral fin wounds, larvae were mounted immediately after wound onto a glass-bottom plate in 1% low melting agarose (Invitrogen) or a custom-built coverslip chamber (for when using an upright scope). Agarose-embedded embryos were covered with 2ml E3 medium (supplemented with tricaine) and imaged either on i) an inverted PerkinElmer UltraVIEW ERS, Olympus IX81 spinning disk confocal microscope with a 30x/1.05 NA silicon (Olympus) or 40x/1.25 NA silicon objective (Olympus) and 488nm for GFP excitation and 561 for tagRFP or mCherry or ii) on an upright Nikon E1000 microscope coupled to a Yokogawa CSU10 spinning disc confocal scanner unit with a 20x/0.75 NA air objective (Nikon) or 10x/0.5 NA air objective (Nikon) and illuminated using a Spectral Applied Research LMM5 laser module (491 nm for GFP excitation; 561 nm for Ruby or TagRFP or mCherry). Confocal stacks using a 2µm z-spacing were acquired every 20-40 sec. Laser wounding was performed on a two-photon scanning miscropscope (LaVision Biotec TriM Scope II). A tunable ultrafast laser (Insight DeepSee, SpectraPhysics) was tuned to 930 nm and the laser power adjusted to approximately 500mW. A square region of interest (ROI) of ∼40µm in width was defined in one focal plane followed by single laser scan across the ROI at a pixel spacing of 240nm and dwell time of 13 µs. Confocal stacks were acquired immediately after, using a 25x/1.05 NA water-dipping lens. GFP was imaged with 930nm and DsRed was imaged with a 1040nm line. For imaging 5-LO translocation, the resolution of imaging with the two-photon microscope was limiting. Larvae were thus transferred (within 10-20 min) for imaging onto an upright Nikon E1000 microscope coupled to a Yokogawa CSU10 spinning disc confocal scanner unit with a 40x/0.80W water objective (Nikon). In some cases, propidium iodide (50µg/ml) was added to the medium 30 min prior to imaging. PI penetration was observed only with superficial laser wound.

For the Cxcl8a response assay, HEK293T cells were cultured in DMEM (Invitrogen) containing 10% FBS (Gibco ThermoFisher Scientific) and 1% Penicillin/Streptomycin (Sigma). HEK293 cells were transfected with Cxcl8-mCherry using Lipofectamine-2000 (Invitrogen) (construct described in (Sarris et al., 2012). Transfected cells were incubated at 37°C (with 5% CO_2_) overnight, harvested the following morning and resuspended in DPBS (Invitrogen) at a density of 30×10^6^/ml. Cells were transplanted above the yolk into 48hpf Tg(*lyz*:Gcamp6F) larvae as previously described (Sarris et al., 2012). Validation of Cxcl8a-mCherry secretion and function *in vivo* is described previously (Sarris et al., 2012).

### Tail fin wounds and Sudan Black staining

The tail fin was amputated up to the level of the notochord and larvae were fixed 20-24 hours later in 1ml of 4% ethanol-free formaldehyde (Polysciences, Warrington, PA) in phosphate-buffered saline (PBS; Sigma-Aldrich) overnight at 4°C with agitation. Fixed larvae were rinsed in PBT (PBS with 0.1% Tween-20; Sigma-Aldrich) twice for 5 minutes and incubated in 1ml Sudan Black (Sigma-Aldrich) for 15 minutes. Following staining, larvae were washed in 70% ethanol for several hours and passaged into 30% ethanol overnight at 4°C with agitation. Larvae were washed in PBT for ten minutes, passaged into increasing concentrations of glycerol and stored in 80% glycerol at 4°C. Larvae were imaged on an optical microscope Stemi 2000-CS (ZEISS) mounted with axiocam ERcs 5s (Zeiss).

### RT-PCR of Cx43 genes in neutrophils

RNA extraction: Larvae were snap frozen in liquid nitrogen after removal of E3 medium. RNA was extracted with the RNAeasy minikit (QIAgen) according to manufacturer’s instruction. RNA was then reverse transcribed with SuperScript™ III Reverse Transcriptase (Invitrogen). PCR was performed using the KOD Hot Start DNA polymerase kit (Novagen, TOYOBO). The list of primers used for RT PCR is indicated on table 3.

**Table 3.**
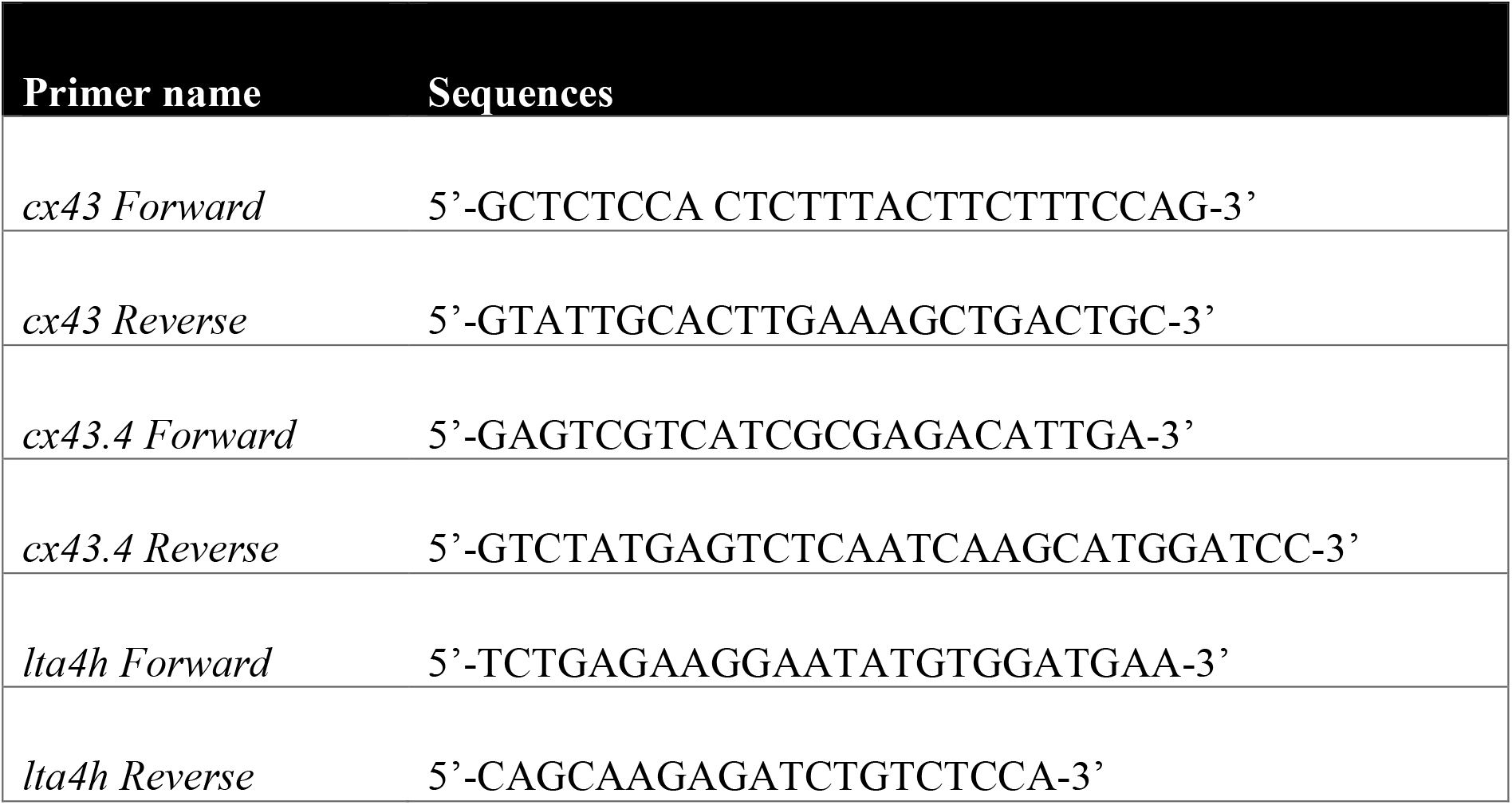
List of primers used for RT PCR.

### Image Analysis

#### Detection and scoring of 5-LO translocation in zebrafish neutrophils

Automated detection of 5-LO tranloscation (Enyedi et al., 2016) was hampered by the irregular shape of the neutrophil nucleus and the fast movement of the cells. We thus used visual inspection of the time-lapse videos on ImageJ (National Institutes of Health, Bethesda, MD). Representative sample videos were confirmed by two viewers and only unambiguous translocation events were scored.

#### Analysis of 5-LO translocation in relation to distance or GCamp6F intensity

Frames in which 5-LO translocation events were detected were duplicated and thereafter analysed with MATLAB 2018b (The MathWorks, Inc., Natick, MA) in an automated fashion. Individual cells were segmented using marker-based watershed segmentation and intensity thresholding. Mean fluorescence intensities of the GCamp6F signal in segmented neutrophils were subsequently computed. For each neutrophil, the fluorescence intensity was normalised to the most fluorescent cell in the corresponding frame to allow pooling of values across embryos with different imaging settings. The wound centre was manually inputted and the distance of individual neutrophil centroids from the wound centre was automatically computed.

#### Analysis of GCamp6F in neutrophil cell-cell contacts

Contacts between bright and dim GCamp6F^+^ neutrophils were counted and categorised according to whether a sharp increase of fluorescence was observed in the dim cell upon contact or not. These events were quantified using visual inspection of the time-lapse videos on ImageJ. Only unambiguous events were scored.

#### Extraction of cell trajectories

Analysis of neutrophil trajectories was performed in Imaris v8.2 (Bitplane AG, Zürich, Switzerland) on 2D maximum intensity projections of the 4D time-lapse videos. Unless otherwise indicated, analysed trajectories were extracted from the area covered by the fin and part of CHT that there was neutrophil immobilisation. A track duration threshold of 3 time-frames was defined to exclude short-lived tracks. Manual track corrections were also applied where needed. Instantaneous (x,y,t) coordinates over time were exported into Microsoft Excel 2016 spreadsheets files (Microsoft Corporation, Redmond, WA). For speed analyses in Figure 5C embryos results are pooled from transgenic Tg(*mpx*:GFP) or Tg(*lyz*:GCamp6F) distributed across the different conditions (no treatment, CBX and morpholino-treated).

#### Extraction of cell surface data

Analysis of neutrophil size and calcium signal was performed in Imaris. Neutrophils were segmented as surfaces and manual surface splitting or merging was applied where needed. Instantaneous neutrophil size and calcium signal intensity were exported into Microsoft Excel 2016 spreadsheets files.

#### Definition of (mechanical or laser) wound

The perimeter of the wound (either mechanical or laser wound) was manually defined as a set of points in MATLAB, using a time-projection of the 2D maximum intensity projection.

#### Quantification of GCamp6F levels

Except for Figure 2 (see previous section on 5-LO-translocation analysis), calcium signal values were extracted from Imaris and imported into MATLAB for plotting. Calcium values for individual segmented neutrophils were normalised to the mean calcium value of the neutrophils in the whole area outside the wound, prior to wounding. For fin wound videos, the calcium values for individual neutrophils were normalised to the mean calcium value of the neutrophils in the whole area outside the wound, at the first timepoint of imaging. For the laser wound experiments, the first 3-9 frames post-wounding were excluded to eliminate distortion of the data by the tissue-scale calcium wave.

#### Quantification of GCamp6F levels with neutrophil size

Neutrophil size and calcium signal values were computed in MATLAB and plotted against each other. A threshold of 100 pixels on the size of detected objects was applied to eliminate false detections.

#### Calculation of neutrophil radial speed

Radial speed was calculated in MATLAB using the following equation (Lämmermann et al., 2013):

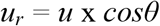

where *u* is the instantaneous speed of neutrophil between two successive positions and *θ* is the angle between the vector of the movement and the vector that connects the position with the wound. The angle *θ* was calculated using the vector between the neutrophil position (centroid) and its nearest point to the wound. When the cosine of *θ* has value 0, the neutrophil migrates directly towards the wound while when it has value −1, the neutrophil migrates directly away from the wound. To decouple trends in directionality of motion from embryo-to-embryo variation in general speed levels, we used a normalisation. Instantaneous speed values for individual neutrophils were divided by the mean instantaneous speed value of the corresponding embryo. Normalised radial speeds were computed with the equation:

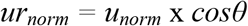

Normalised radial speed values were binned every 7.5 minutes. For the laser wound experiments, the first 3-9 frames post-wounding were excluded for consistency with the calcium signal calculation.

#### Quantification of neutrophil GCamp6F levels and speed upon contact with necrotic cells

Individual neutrophils were visually inspected to determine the time-point at which they touched the PI-stained necrotic cells. This time-point was considered as the time-point 0. The neutrophils were tracked for 180 seconds before and after this time-point. Individual neutrophil calcium values were normalised with the calcium value of the first time-point of the track.

##### Statistics

All error bars indicate S.E.M. All p values were calculated with two-tailed statistical tests and 95% confidence intervals. t-test (pairwise comparisons) and one-way ANOVA (multiple group comparisons) were performed after distribution was tested for normality otherwise non-parametric tests were performed (Mann-Whitney for two-way comparisons and Kruskal-Wallis with Dunn’s post-test for multiple comparisons). Statistical tests were performed in Prism8 (GraphPad Software Inc., La Jolla, CA). The statistical test and the n number are indicated in the figure legends. The error bars show standard error of the mean across individual embryos (reflecting embryo variation), except for the descriptive statistics in Figure 3C, 5C and 5E, where error bars reflect variation across cells pooled from different embryos. Live imaging experiments were acquired in minimum three independent experiments. In figure panels, * corresponds to p<0.03, ** to p<0.002 and *** to p<0.0002.

