## Supplemental Figures 1-6 and Video Legends for "Neutrophil swarming in damaged tissue is orchestrated by connexin-dependent calcium signals"

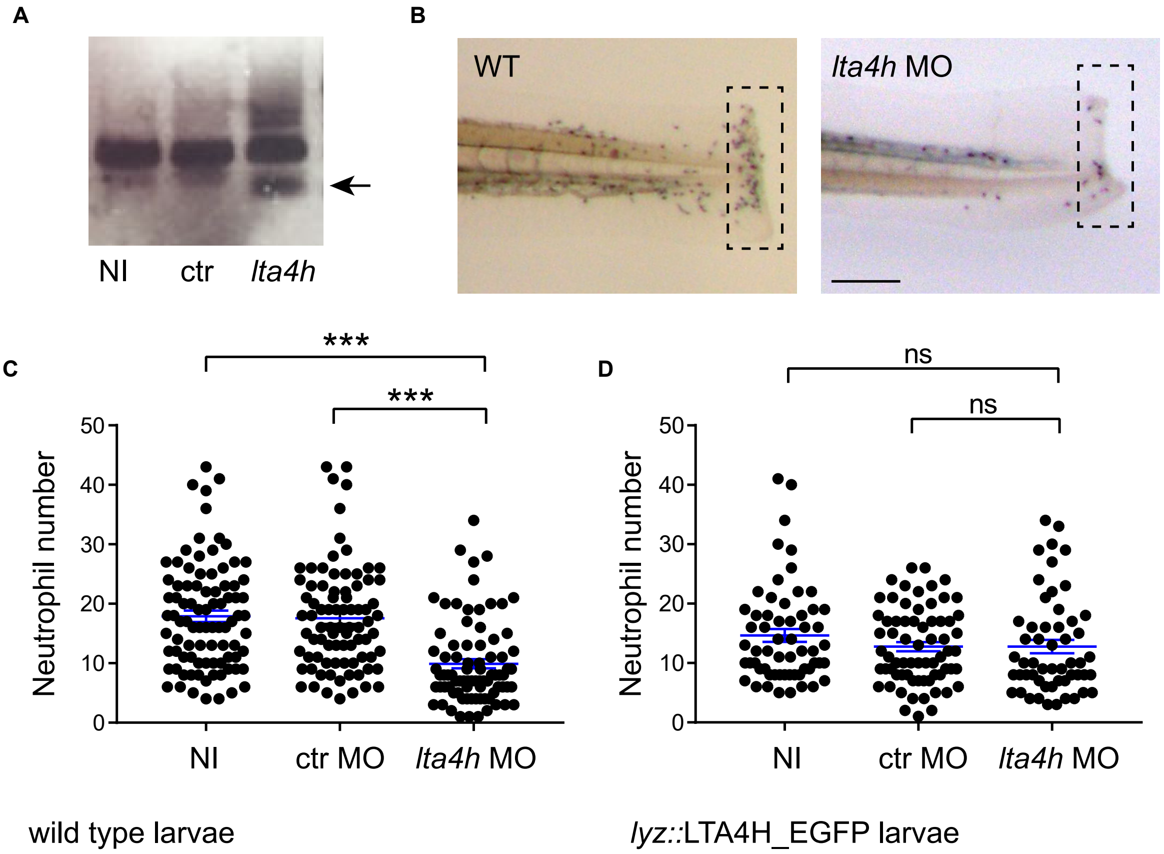


**Figure S1. Role of neutrophil biosynthesis of LTB4 in zebrafish wounds, Related to Figure 1 and 2A.**

**A** RT-PCR of *lta4h* from mRNA extracted from 3 dpf zebrafish. The arrow denotes the presence of an alternative transcript in larvae injected with a splice-blocking *lta4h* morpholino; NI = Non-Injected, ctr = Control, *lta4h* = injected with the splice-blocking *lta4h* morpholino.

**B** Sudan black staining of neutrophils in WT and *lta4h* morpholino-injected 3dpf larvae. Embryos were injured with a scalpel and fixed 3h after wounding. The dotted lines represent the area in which neutrophils were counted; scale bar = 100µm.

**C**, **D** Quantification of the total number of neutrophils recruited to control and morphant wounds.

**C** 81-89 larvae per group pooled from 3 experiments.

**D** 54-68 larvae per group pooled from 3 different experiments. p value <0.0001. Kruskal-Wallis test with Dunn's post-test.

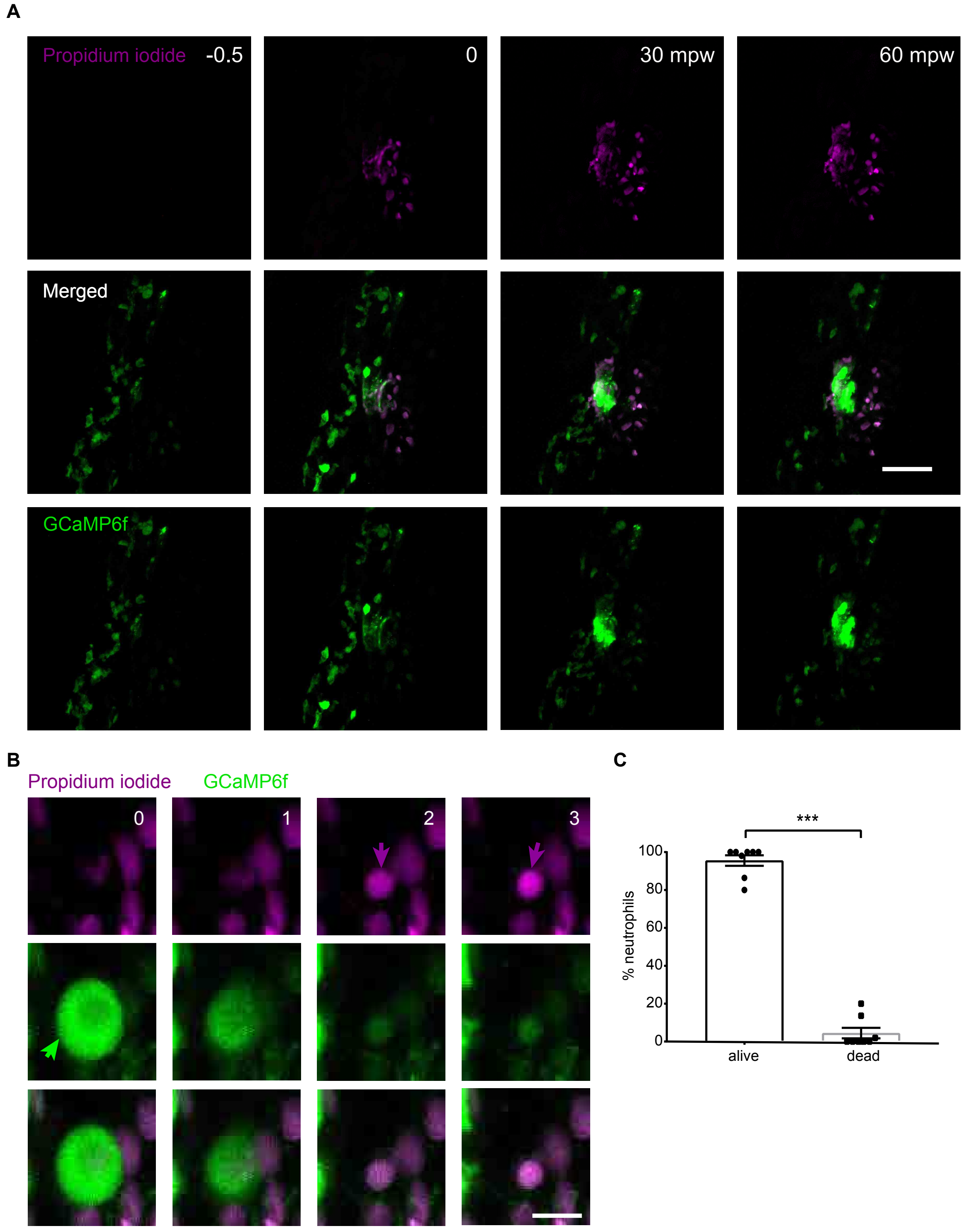


**Figure S2. Percentage of dying cells among neutrophils clustering at the wound, Related to Figure 3.**

**A** Confocal time-lapse projections of neutrophils expressing Gcamp6F (green) in 3dpf larvae incubated in PI. Dotted line indicates laser wound.

**B** Zoomed in example of an apoptotic neutrophil. The green arrow indicates a neutrophil with an apoptotic shape in the GCamp6F channel. The red arrows indicate the nucleus of this neutrophil subsequently up taking PI. Scale bar = 10µm.

**C** Percentage of dead or alive neutrophils averaged through the first hour post-wound. Data pooled from 8 larvae in 7 independent experiments. Mann-Whitney test, two tailed, p-value= 0.0002.

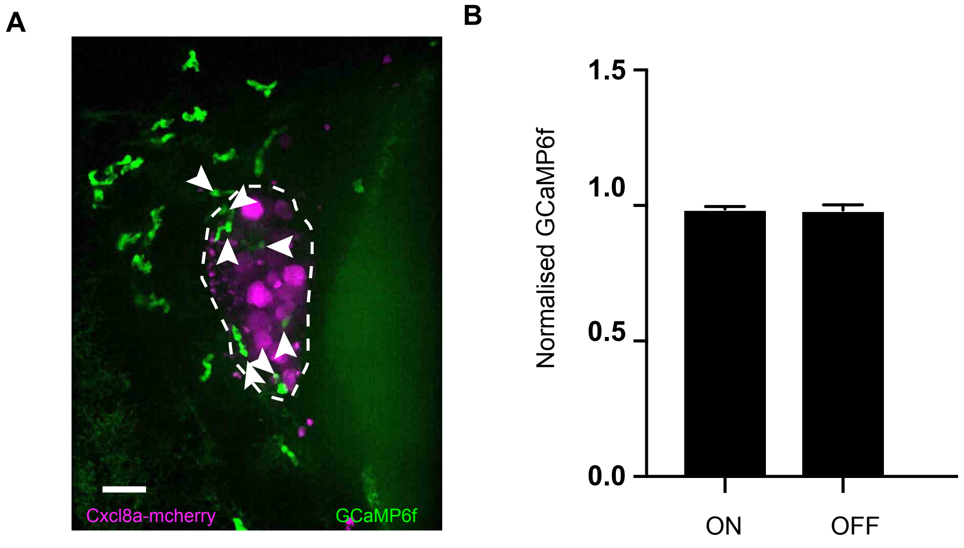


**Figure S3. A cellular chemokine source does not trigger a calcium alarm signal in neutrophils, Related to figure 4.**

**A** Confocal spinning disk projections of neutrophils in Tg(*lyz*:Gcamp6f) 3 dpf larvae (in green) moving around a transplant of HEK293T cells expressing Cxcl8a-mCherry (in red). The transplantation zone is delimited by dotted lines and neutrophils within this area are indicated with white arrows. This image corresponds to Supplemental video 10. Scale bar = 25µm. See also video S8.

**B** Quantification of GCamp6F intensity in neutrophils on the Cxcl8a transplant (ON) and beyond (OFF). Data are means of 4 independent embryos.

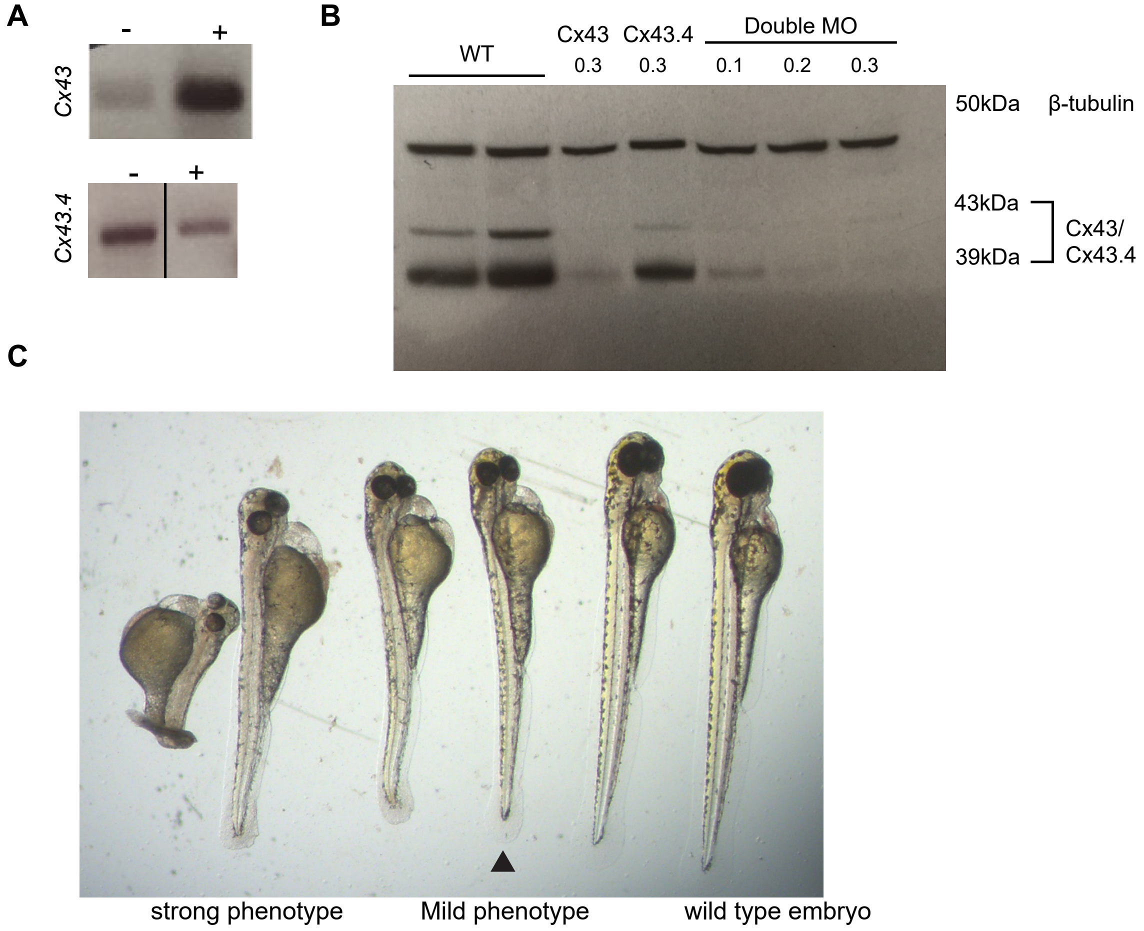


**Figure S4. Neutrophil Cx43 expression and knockdown in zebrafish larvae, Related to Figure 5.**

**A** RT-PCR for Cx43 and Cx43.4 expression from cDNA samples of FACS-sorted GFP$^+$ (+) or GFP$^-$ cells (-) from Tg(*mpx*:GFP) 4.5 dpf larvae.

**B** Western Blot showing the expression of Cx43 and Cx43.4 in 3dpf larvae in WT and morphant zebrafish larvae. Amount injected for each morpholino in pmol is shown.

**C** Image of different phenotypes of 3dpf zebrafish larvae injected with cx43/cx3.4 combination MOs. The black arrow indicates the mildest phenotype with detectable difference in eye size that was selected for subsequent experiments.

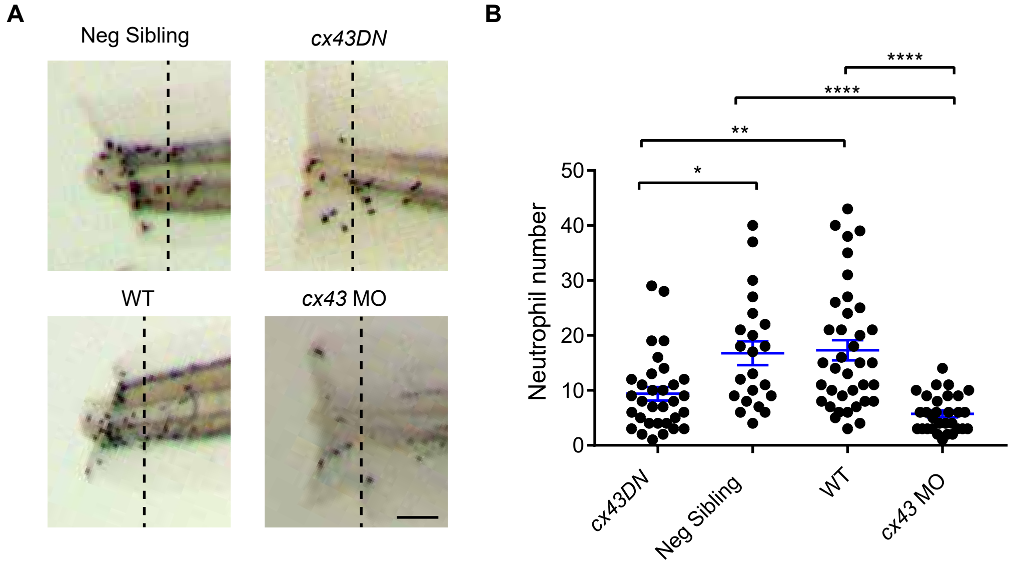


**Figure S5: Cx43 inhibition leads to reduced neutrophil accumulation at mechanical fin wounds, Related to Figure 5.**

**A** Sudan black staining of neutrophils in transgenic Tg(*lyz:cx43DN*-T2A-mCherry) larvae (*cx43DN*) and their negatively screened siblings (sibling) WT larvae, larvae injected with (*cx43/cx43.4 combination of* morpholinos (*cx43* MO). 3 dpf larvae were amputated with a scalpel and fixed 3h after wounding. The dotted lines represent the area in which neutrophils were counted. Scale bar = 50µm.

**B** Quantification of the total number of neutrophils recruited to wounds in the different conditions pooled from three independent experiments. *dn cx43,* n= 32 larvae; negative siblings, n=22 larvae; wt, n=37 larvae; *cx43* MO, n= 30 larvae. Kruskal-Wallis multiple comparisons test with Dunn's post-test.

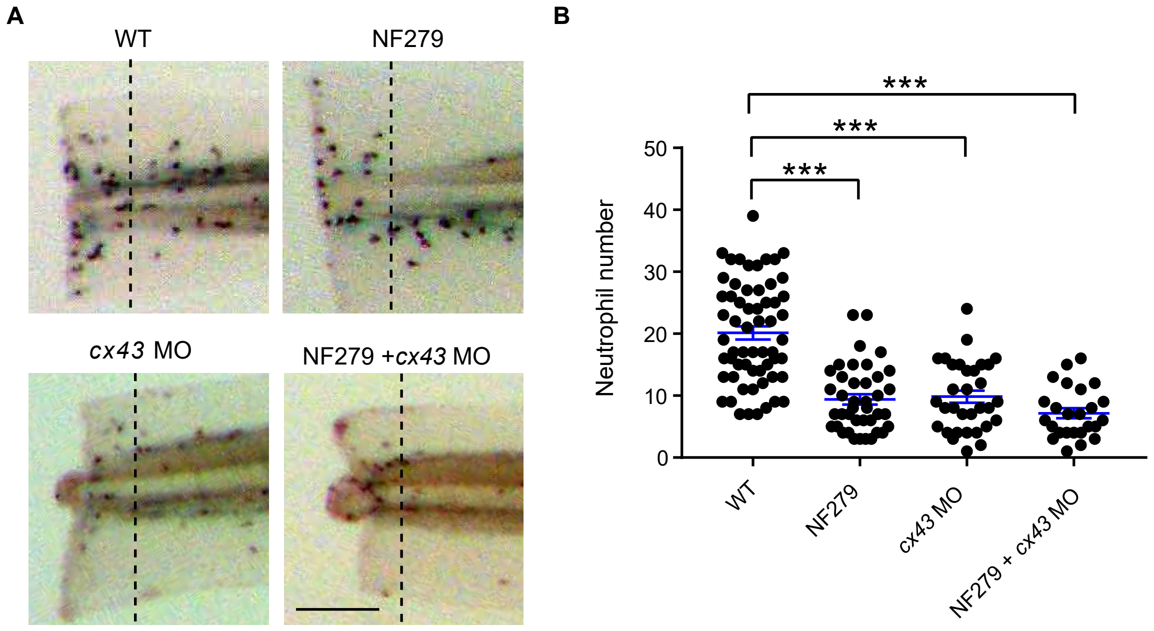


**Figure S6. Absence of additive effect of Cx43 inhibition on neutrophil migration over P2X1 inhibition, Related to Figure 4 and 5.**

**A** Sudan black staining of neutrophils in wild type larvae injected or not with *cx43/cx43.4 combination of* morpholinos (*cx43* MO) and treated or not with 10µM NF279. 3 dpf larvae were amputated with a scalpel and fixed 3h after wounding. Scale bar =100µm.

**B** Quantification of neutrophil number at wounds in the different conditions. Data pooled from two independent experiments. One-way ANOVA with Tukey's multiple comparison test. WT (non-injected and not treated with drug), n=60 larvae, NF279, n=40, cx43 MO n=32, cx43 MO with NF279 n=25.

### Supplemental Video Legends

**Video S1. Distinct calcium signaling patterns in clustering neutrophils,** (Subvideo 1a) Neutrophils in Tg(*lyz* :Gcamp6f) larvae responding to a laser-induced focal tissue damage in the caudal hematopoietic tissue-ventral fin boundary. The same video is shown either colour-coded for intensity (left), or grayscale at high (middle) and low brightness (right) to enable visualisation of calcium dynamics in migrating and clustering cells respectively. The analysis of the clustering response is presented in Figure 1c and Figure 1d. Maximum intensity projections from two-photon microscopy z stacks are shown. Scale bar= 25µm. Frame interval is 30 sec. (Subvideo 1b) Calcium dynamics in small clusters. A similar example as subvideo 1a but with small-scale recruitment. Acquisition details are the same. Scale bar = 50µm. See Figure 1B.

**Video S2. Neutrophil contacts propagate calcium fluxes.**
Series of examples of neutrophils in different Tg(*lyz* :Gcamp6f) larvae showing propagation of the calcium signal from one cell to another. Neutrophils with low calcium levels encountering neutrophils with high calcium levels are shown with an arrow. Time is indicated in minutes. Image dimensions in µm in x,y: Stack1 42x32; Stack2 40x30; Stack3 79x67; Stack4 53x56 ; Stack 5 25x26 ; Stack 6 137x103. See Figure 3D.

**Video S3. 5-LO translocation in neutrophils at laser wounds and mechanical wounds.** (subvideo a) Neutrophils in Tg(*lyz*:Gcamp6f) (left) crossed with Tg(*lyz*:5LO-tRFP) (right) responding to a laser wound damage. One of the 5LO-tRFP translocation events is highlighted in slow motion. The analysis of the 5LO-tRFP translocation is presented in

Figure 2. Two-photon microscopy was used for wounding and spinning disc microscopy for imaging. Scale bar = 15µm. Time-lapse every 30sec over 60 min (video plays at 15 frames/s). (subvideo b) 5-LO translocation in neutrophils at mechanical fin wounds. Neutrophils in Tg(*lyz*:Gcamp6f) (left) crossed with Tg(*lyz*:5LO-tRFP) (right) responding to a ventral fin wound. One of the 5LO-tRFP translocation events is highlighted in slow motion. Imaging starts 10 min post wound (note that we miss the tissue calcium wave as imaging does not start immediately after wounding). The analysis of the 5LO-tRFP translocation is presented in Figure 2. Spinning disc microscopy was used approximately 15 min post mechanical wound. Scale bar = 15µm. Time-lapse every 1 min over 135 min (video plays 15 frames/s). See Figure, 2C.

**Video S4. Neutrophil contact with necrotic tissue triggers calcium elevation.** Neutrophils in Tg(*lyz*:Gcamp6f) larvae incubated in PI (red) responding to a two-photon laser wound tissue damage. A neutrophil contact with $PI^+$ necrotic cells is highlighted during the video. The corresponding analysis is presented in Figure 3. Maximum intensity projection of z-stacks from two-photon microscopy is shown. Scale bar= 25µm. Time-lapse every 30sec over 75 min 30sec (video plays at 15 frames/s). See Figure 3B.

**Video S5. Dying neutrophils ejected from the cluster.**

Series of examples of neutrophils in different Tg(*lyz*:Gcamp6f) larvae, two of which incubated with PI (red) (subvideos 2 and 3 in the sequence), showing neutrophil death/apoptosis and ejection from the cluster. Dying neutrophils are indicated with white arrows. Time is indicated in minutes. Image dimensions in µm in x,y: Stack1 53x51; Stack2 59x59; Stack3 49x47; Stack4 46x51. See Figure S2B, C and D.

**Video S6. Neutrophil migration and calcium dynamics in a ventral fin wound in the presence or absence of NF279**.

Neutrophils in Tg(*lyz*:Gcamp6f) larvae incubated with NF279 (right) or in control non-treated larvae (CTR, left) responding to a mechanical fin wound. Imaging starts 10 min post wound. Maximum intensity projection of a z-stack is shown. Scale bar= 25µm. Time-lapse is every 30sec over 120 min (video plays at 15 frames/s). Image dimensions in µm in x, y: control 207x277; NF279 275x367. See Figure 4B.

**Video S7. Neutrophil calcium dynamics upon chemokine stimulation.** Neutrophils in the head of 3 dpf Tg(*lyz*:Gcamp6f) (green) larvae responding to a transplant of Cxcl8a-mCherry-expressing HEK293T cells (red). Spinning disc microscopy was used. Maximum intensity projection of a z-stack is shown. Scale bar= 25µm. Time-lapse is every 20 sec over 40 min (video plays at 15 frames/s). See Figure S3A.

**Video S8. Extracellular calcium entry is triggering stopping and 5-LO translocation in neutrophils.** Neutrophils in Tg(*lyz*:Gcamp6f) (left) crossed with Tg(*lyz*:5LO-tRFP) (right) responding to a ventral fin wound. 50µM calcium ionophore is added 45min into the video. Spinning disc microscopy was used. Scale bar= 25µm. Time-lapse is every 30sec over 84 min (video plays at 15 frames/s). See Figure 4F.

**Video S9. Calcium pattern and neutrophil behaviour in the presence of CBX or upon treatment with *cx43* MO.** (subvideo 1) Neutrophils in Tg(*lyz*:Gcamp6f) larvae incubated in PI (red) responding to a two-photon laser wound tissue damage in the presence of 50µM Carbenoxolone. Scale bar= 25µm. Note that the initial tissue calcium wave is present but not the fluxes in neutrophils at the wound. Two-photon microscopy was used. Max intensity projection of a z-stack is shown. Time-lapse is every 30 sec over 149.5 min (video plays at 15 frames/s). (subvideo 2) Neutrophils in Tg(*lyz*:Gcamp6f) larvae injected with a combination of *cx43* MOs and incubated in PI (red) responding to a two-photon laser wound tissue damage. Note that the initial tissue calcium wave is present but not the fluxes in neutrophils at the wound. Maximum intensity projection of z-stacks from two-photon microscopy are shown. Scale bar=50µm. Time-lapse every 30sec over 87.5 min (video plays at 15 frames/s). See Figure 5A-C.

**Video S10. Calcium signalling pattern and neutrophil behaviour at wounds after neutrophil-specific inhibition of Cx43.** Neutrophils in Tg(*lyz*:Gcamp6f)xTg(*lyz*:*cx43*DN_T2A-mCherry) larvae positive for the *cx43*DN_T2A-mCherry construct (right) or not (control sibling - left) responding to a mechanical fin wound. Imaging starts 10 min post wound. Scale bar= 25µm. Maximum intensity projection of a z-stack is shown. Time-lapse is every 30sec over 149.5 min (video plays at 15 frames/s). See Figure 5D-E.

**Video S11. Defective propagation of calcium signals in neutrophils after *cx43* inhibition.** Series of examples of neutrophils in Tg(*lyz*:Gcamp6f) larvae showing propagation of the calcium signal from one cell to another in different conditions: cross with TgxTg(*lyz*:*cx43*DN_T2A-mCherry), *cx43* morpholino injection and CBX treatment. Neutrophils with low calcium levels coming into contact with neutrophils with high calcium levels are shown with an arrow. Time is indicated in minutes. Image dimensions in µm in x,y: Stack1 36x27; Stack2 24x30; Stack3 38x34; Stack4 75x75; Stack5 61x62. See Figure 5F-I.
